# Gram-negative Antimicrobial spectrum of *Ulmus pumila*

**DOI:** 10.1101/2022.03.12.484112

**Authors:** Josh Schafer, Troy B. Puga, Sha’Rai T. Miller, Prince N. Agbedanu

**Affiliations:** University of Kansas School of Medicine, Kansas City, USA; Kansas City University School of Medicine, Kansas City, USA; Hutchinson Community College, Newton, Kansas, USA; Friends University, Department of Health Sciences, Wichita, Kansas, USA

**Keywords:** antimicrobial, bacteria, gram-negative, drug resistance, antimicrobial spectrum, microbiology

## Abstract

**Purpose:** This study explores the antimicrobial spectrum of *Ulmus pumila* by screening commonly infectious gram-negative bacteria for susceptibility. Currently, many gram-negative bacteria are developing resistance to recommended antibiotic therapies used in clinical practice. It is imperative that we continue to explore new antimicrobial compounds in order to help combat the global health issue of antibiotic resistance.

**Method:** 3g of freshly harvested leaves and flower samples of *Ulmus pumila* were extracted in 15 ml of 95% ethanol. The mixture was filtered to remove the debris and sterile blank discs were soaked in the clear filtrate (samples) or the extraction solvent (as background controls) for 20 minutes. Glycerol stocks of bacteria were scaled in LB broth and Muller Hinton agar was prepared with 38g of agar in a liter of water. A 100-microliter suspension of the scaled bacteria was diluted with 9 ml of 1% saline solution and 100 microliters of this saline dilution was plated. Sterile paper discs that were infused with extracts (samples) or vehicle control (95% ethanol) were placed on the freshly plated bacterial plates and incubated at 37 degree Celsius overnight. Zones of inhibition were recorded as a measure of antibacterial activity.

**Result:** *Ulmus pumila* demonstrated antimicrobial activity against the following gram-negative bacteria: *E. coli* (16 mm mean zone of clearing), *P. vulgaris* (15 mm mean zone of clearing), *E. cloacae* (20.5 mm mean zone of clearing), and *K. pneumoniae* (16 mm mean zone of clearing).

**Conclusion:** *Ulmus pumila* displayed antimicrobial activity against various species of gram-negative bacteria including *E. coli, P. vulgaris, K. pneumonaie, and E. cloacae. Ulmus pumila* has also previously demonstrated activity against some species of gram-positive bacteria. By this discovery, *Ulmus pumila* has the potential to serve as a broad-spectrum antimicrobial agent with activity against both gram-positive and gram-negative bacteria.

## Introduction

Drug-resistant strains of bacteria are a growing concern for clinicians and public health experts across the globe. Antibiotic resistance has cost the healthcare system, as a whole, billions of dollars [1,2]. Antibiotic resistance has also led to many deaths and is likely to worsen across the globe [1,2]. Failure to develop new antibiotics has greatly contributed to antibiotic resistance [1]. This research investigates the antimicrobial properties of *Ulmus pumila* against gram-negative bacteria so that it may help combat the growing issue of drug resistance.

*Ulmus pumila*, or Siberian Elm, is a fast growing and invasive species that is native to Northern China, Eastern Siberia, Manchuria, and Korea, but also found throughout the United States [3]. It is considered invasive in 23 states, and can grow up to 70 feet in height [3]. It has previously been found that *Ulmus pumila* may have activity against *methicillin-resistant Staphylococcus aureus* (MRSA) and also may have cytotoxic properties that make it useful as an anticancer therapy [4,5]. This experiment tests the activity of *Ulmus pumila* against the following pathogenic gram-negative organisms: *E. coli, P. vulgaris, K, pneumoniae, and E. cloacae*.

Gram-negative bacteria are highly drug-resistant bacteria and are considered more resistant than gram-positive bacteria [6]. Gram-negative bacteria are a significant cause of morbidity and mortality worldwide due to the ability of gram-negative bacteria to cause infections in almost all human organ systems [6, 7]. Drug-resistant gram-negative bacteria significantly increase the burden in the ICU [8]. Drug-resistant gram-negative bacteria are responsible for most cases of ventilator-associated pneumonia and other ICU-acquired sepsis pathogens [6]. Gram negative bacteria can develop drug-resistance through several mechanisms such as efflux pumps, alterations in membrane permeability, and degradation enzymes, which means the bacteria are always evolving [7]. Development of new antibiotics is critical to fight growing drug resistance by gram-negative bacteria [6,7].

This experiment tests the antimicrobial properties of *Ulmus pumila* against the gram-negative *E. coli, P. vulgaris, K, pneumoniae, and E. cloacae. Escherichia coli (E. coli*) is one of the most common bacterial infections in humans [9]. *E. coli* is a gram-negative bacterium that exists throughout the gastrointestinal tract [10]. *E. coli* is a common cause of urinary tract infections, bacteremia, and neonatal meningitis in humans [9,10]. *E. coli* has several pathogenic strains that can cause diarrheal illness and vomiting [9,10]. One particular strain associated with watery diarrhea is ETEC (Enterotoxigenic *E. coli*), also known as travelers’ diarrhea [9,10]. A particular strain of bloody diarrheal illness is a subtype of EHEC (Enterohaemorrhagic *E. coli)* called *E. coli* O157:H7, which can cause a potentially lethal infection in children called Hemolytic Uremic Syndrome [9-11]. Most notably, *E. coli* has demonstrated increased resistance and decreased susceptibility to many strains of antibiotics [9]. *Proteus vulgaris* is a gram-negative rod that is part of the normal flora, but has been known to cause community and hospital acquired urinary tract infections [12,13, 14]. *Proteus vulgaris* has demonstrated drug resistance against several antibiotics used in clinical practice [15].

*Klebsiella pneumoniae (K. pneumonaie)* is a gram-negative bacterium that colonizes the oropharynx and gastrointestinal tract [16]. *K. pneumoniae* is the most common cause of nosocomial pneumonia in the United States, and is also a common cause of urinary tract infections and bacteremia [16, 17]. *K. pneumoniae* has developed antibiotic resistance against many antibiotics, and is becoming a very big concern for healthcare experts around the world [17,18]. The final gram-negative pathogen tested in this experiment is *Enterobacter cloacae (E. cloacae). E. cloacae* is a gram-negative rod that is part of humans’ natural flora of the gut, but has been associated with large nosocomial infection outbreaks [19]. *E. cloacae* has been known to cause bacteremia, endocarditis, and septic arthritis [19]. *E. cloacae* is becoming a major global threat due to its drug resistance against last-resort antibiotics [19,20].

The purpose of this study is to investigate the antimicrobial properties of *Ulmus pumila* against several pathogenic species of gram-negative bacteria for potential development as a broad-spectrum antimicrobial. We hypothesize that *Ulmus pumila* will demonstrate antimicrobial activity against the gram-negative bacteria *E. coli, P. vulgaris, K. pneumoniae*, and *E. cloacae*.

## Method

### Sample extraction, disc preparation and plating

3g of freshly harvested leaves and flower samples of *Ulmus pumila* were homogenized with a mortar and pestle. The homogenate was suspended in 15 ml of 95 % ethanol [21] and swirled for 15 minutes to allow time for extraction of active ingredients. The mixture was filtered to remove the debris. Sterile blank discs were soaked in the clear filtrate containing the active ingredient for 20 minutes as previously described [21]. The extraction solvent (95% ethanol) was also infused into a blank disc to control for vehicle effect. Glycerol stocks of bacteria were scaled as described previously [21] and agar plates were also prepared as described previously [21]. One hundred microliter suspension of the scaled bacteria was diluted with 9 ml of 1% saline solution and 100 microliters of this dilution was plated [21]. Discs that were infused with extracts or vehicle control (95% ethanol) were placed on bacterial plated plates. The plates were incubated at 37 degree Celsius overnight and zones of inhibition were measured.

## Result

*Ulmus Pumila* showed antimicrobial activity against the following gram-negative bacteria: *E. coli* (16 mm mean zone of clearing), *P. vulgaris* (15 mm mean zone of clearing), *E. cloacae* (20.5 mm mean zone of clearing), and *K. pneumoniae* (16 mm mean zone of clearing).

## Discussion

The results of this study demonstrated *Ulmus pumila* has antimicrobial activity against the following gram-negative bacteria: *E. coli, K. pneumoniae, E. cloacae*, and *P. vulgaris*. These bacteria are of concern as they have all been implicated in causing severe infections and showing drug-resistance to common antibiotics used in clinical practice [9-20]. *U. pumila* has also been proven to show modest gram-positive coverage and potential anti-cancer effects [4,5]. Antibiotic resistance is a significant cause of mortality worldwide and has demonstrated significant financial strain on health care systems, making the discovery and development of novel antibiotics of vital importance [1,2]. Gram-negative bacteria are a particular concern in regards to their high levels of drug resistance [6]. With the results of this study and previous studies, we conclude that *Ulmus pumila* has the potential to be developed as a broad-spectrum antibiotic due to its antimicrobial properties against gram-negative and gram-positive bacteria.

We acknowledge that this study is limited due to the inability to test against drug-resistant strains of gram-negative bacteria This limitation was due to biohazard concerns of the facility. We encourage researchers with secure facilities to test drug-resistant strains against *U. pumila*. We believe this research serves as an excellent first step in the development of further research into *U. pumila*. Further research experiments of *U. pumila* may include testing drug-resistant bacterial strains, testing for side effect profiles, and testing for antifungal properties. We encourage researchers to pursue these potential research opportunities.

## Acknowledgment

We thank the VPAA’s office and the Chair of STEM, Dr. Nora Strasser, of Friends University, for their support of both undergraduate and graduate research. We thank our collaborator, Mr. Nasir Islam for his help on the maintenance of bacterial cultures.

## Author Contribution

Conceptualization, PNA; Data curation, TP; Investigation, STM and PNA; Methodology, PNA; Writing – original draft, JS and TP; Visualization, JS; Writing – review & editing, JS and PNA

---

**Figure 2:**
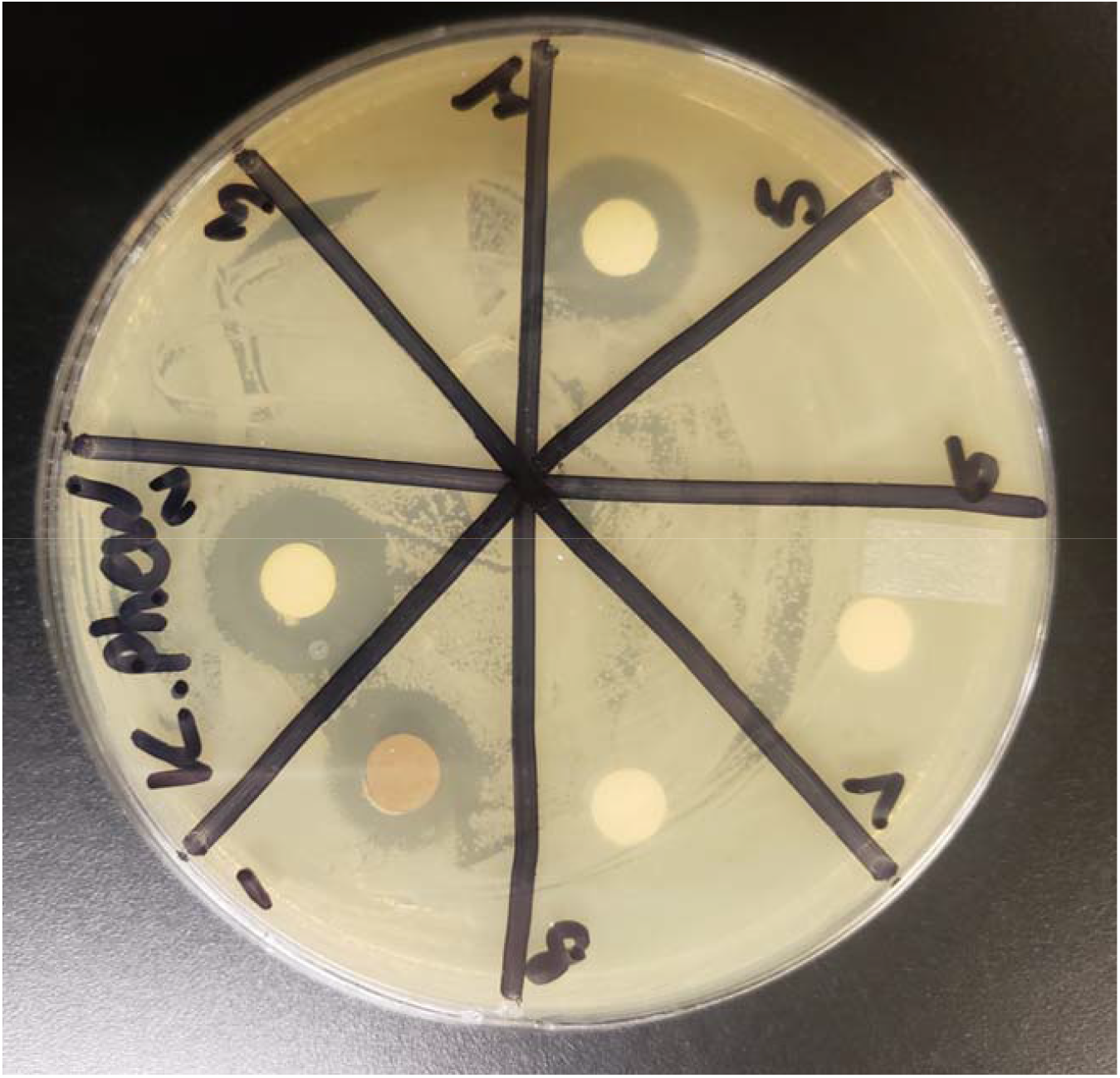
*Ulmus pumila* (Position 2) and blank disk (Position 8) against *K. pneumoniae*.

**Table 1:** Gram negative bacteria zone of clearing (in mm) against a blank disk and *Ulmus pumila*.

|  | *E. coli*<br>(clearing in mm) | *P. vulgaris*<br>(clearing in mm) | *E. cloacae*<br>(clearing in mm) | *K. pneumoniae*<br>(clearing in mm) |
| --- | --- | --- | --- | --- |
| Blank Disk | 0 mm | 0 mm | 0 mm | 0 mm |
| *Ulmus pumila* | 16 mm | 15 mm | 20.5 mm | 16 mm |

**Figure 2:**
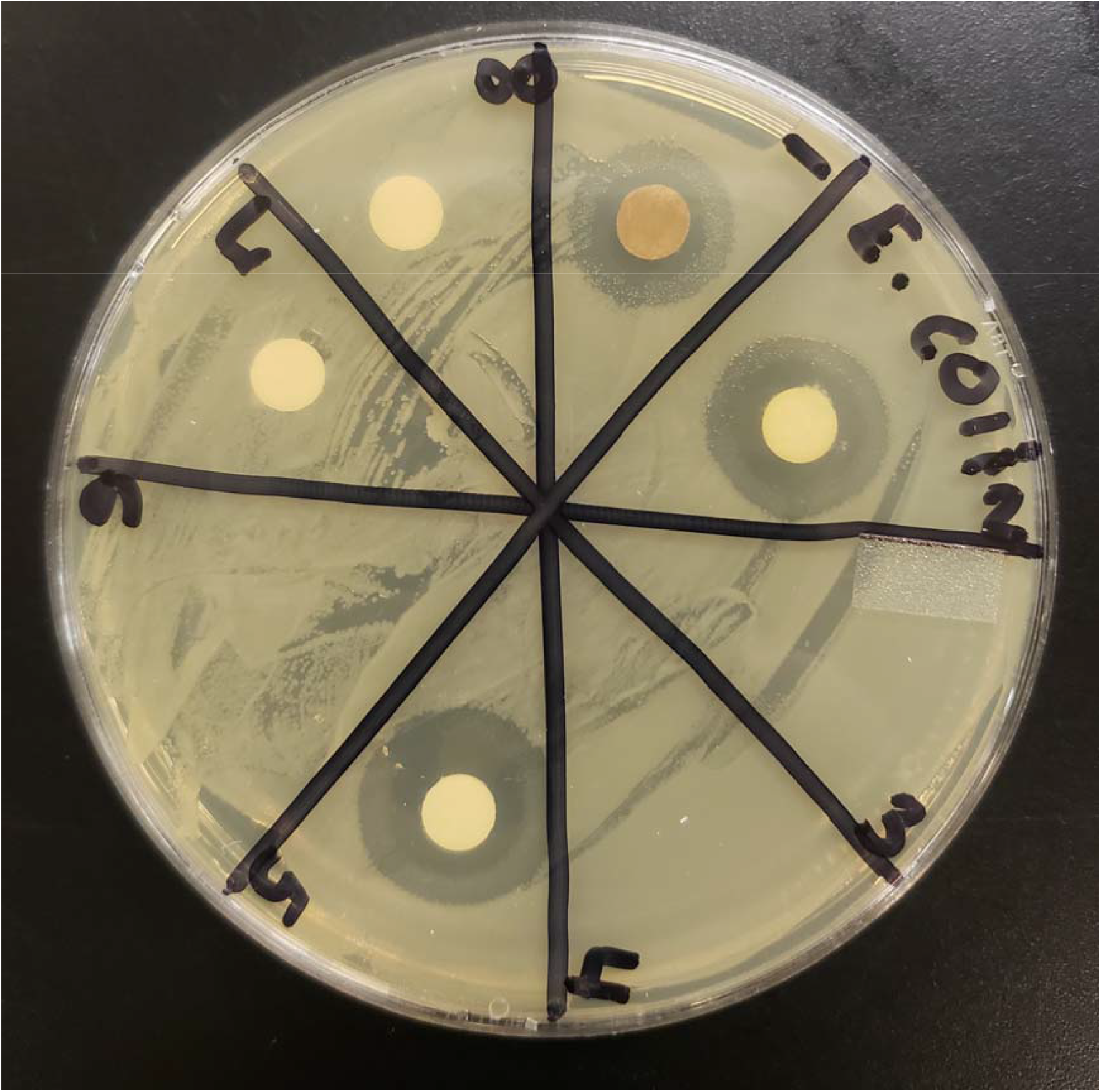
*Ulmus pumila* (Position 2) and blank disk (Position 8) against *E. coli*.

